# Using in silico perturbational approach to identify critical areas in schizophrenia

**DOI:** 10.1101/2022.12.15.520260

**Authors:** Ludovica Mana, Manel Vila − Vidal, Charlotte Köckeritz, Kevin Aquino, Alex Fornito, Morten L. Kringelbach, Gustavo Deco

## Abstract

Schizophrenia is a debilitating neuropsychiatric disorder whose underlying correlates remain unclear despite decades of neuroimaging investigation. One contentious topic concerns the role of global signal fluctuations and how they affect more focal functional changes. Moreover, it has been difficult to pinpoint causal mechanisms of circuit disruption. Here we analysed resting-state fMRI data from 47 schizophrenia patients and 118 age-matched healthy controls and used dynamical analyses to investigate how global fluctuations and other functional metastable states are affected by this disorder. We then used in-silico perturbation of a whole-brain model to identify critical areas involved in the disease. We found that brain dynamics in the schizophrenic group were characterised by an increased probability of globally coherent states and reduced recurrence of a substate dominated by coupled activity in the default mode and limbic networks. Perturbing a set of temporoparietal sensory and associative areas in a model of the healthy brain reproduced global pathological dynamics. Healthy brain dynamics were instead restored by perturbing a set of medial fronto-temporal and cingulate regions in the model of pathology. These results highlight the relevance of global signal alterations in schizophrenia and identify a set of vulnerable areas involved in determining a shift in brain state.

## 1. INTRODUCTION

Schizophrenia is a debilitating condition that imposes a heavy burden for both patients and society. Characterised by a great heterogeneity of manifestations and associated with a diverse array of positive, negative, and cognitive symptoms, the disorder is thought to have a neurodevelopmental origin, affects numerous brain regions, and is characterised by variable disease progression and response to treatment (Mueser and McGurk 2004).

Schizophrenia has long been proven to be directly associated with brain dysfunction, and plenty of studies have been looking for biological markers. In particular, extensive neuroimaging research using techniques such as functional magnetic resonance imaging (fMRI) and diffusion-weighted imaging (DWI) tractography has been carried on in the past few decades, and has provided evidence of widespread structural and functional MRI brain alterations in psychosis (Gur and Gur 2010; Kraguljac et al. 2021). However, many results are contradictory and only partially replicable, especially when considering alterations of brain functional connectivity (FC) (Fitzsimmons, Kubicki, and Shenton 2013; Fornito et al. 2012; Fornito and Bullmore 2015). While a consistent portion of literature describes local and global reductions of functional connectivity, consistent with repeated observations of reduced structural connectivity (Friston and Frith 1995; Pettersson-Yeo et al. 2011), some works report increased level of FC instead (Chopra et al. 2021; Jafri et al. 2008; Whitfield-Gabrieli et al. 2009; Yang et al. 2014).

Such inconsistencies may be due to high heterogeneity of the patients diagnosed with this disorder (Cole et al. 2011; Feczko et al. 2019; Wolfers et al. 2018), and potentially by an inadequate conceptualisation of the disease itself (Deacon and McKay 2015; Marquand et al. 2019). However, the problem can also be partially explained by methodological issues such as an emphasis on studying time-averaged, static FC when functional changes in brain activity are intrinsically dynamical and time-varying. Various methods have been developed to capture dynamical properties of resting-state fMRI data beyond classical static functional connectivity, among which LEiDA analysis (Leading Eigenvector Dynamics Analysis) has been repeatedly proven to effectively characterise differences in a number of different conditions (Cabral et al. 2017; Escrichs et al. 2022). Another controversial issue regards the debate on pre-processing in fMRI data; in particular the problems posed by the interpretation of globally coherent signal fluctuations. Such fluctuations, often quantified as the mean signal averaged across the entire brain and referred to as the global signal, have long been considered to be noise, and therefore generally removed from the data using a procedure called global signal regression (GSR) (Aquino et al. 2020; Liu, Nalci, and Falahpour 2017; Power et al. 2017). Several studies though have highlighted how correction of this signal may alter some of the effects of interest(Glasser et al. 2019; Zhang and Northoff 2022). In this regards, some papers pointed out the importance of deeply investigating its role in schizophrenia (Yang et al. 2014, 2017).

Finally, it is important to note that very few findings identified in case-control comparisons of FC have proven clinically actionable, mainly due to a difficulty in determining how any identified FC differences directly relate to the disease (Swerdlow 2011). This lack of clinical applicability underscores the need to develop novel strategies to gain insights into the causal role of such alterations. Even though several mechanisms have been proposed (Rolls et al. 2008), it is difficult to fully reproduce the complex combination of clinical symptoms of schizophrenia in animal models, where direct manipulation is possible (Marcotte, Pearson, and Srivastava 2001; Young, Zhou, and Geyer 2010), slowing down the clinical translation of research findings. Computational models offer a bridge between measured imaging signals and underlying cellular and molecular mechanisms, allow to investigate the complex interaction between functional and structural properties of the data (Huys, Maia, and Frank 2016; Krystal et al. 2017), and provide an opportunity for testing different models of causal mechanisms and to infer direct relation by mean of perturbational approaches (Deco et al. 2019; Deco and Kringelbach 2014; Deco, Vidaurre, and Kringelbach 2021).

Here, we investigated dynamical changes in the functional organisation of the brain in people with schizophrenia. In particular, we applied dimensionality reduction and clustering approach on a measure of dynamical functional connectivity, to accurately separate the alterations related to global signal from alterations in a specific pattern of activation. We then used *in silico* perturbation of a whole-brain model to identify areas involved in the emergence of the disease and in potential compensatory mechanisms.

To do so, we adapted a pipeline recently developed by our group (Deco et al. 2019) which effectively detects critical areas involved in different altered states (Escrichs et al. 2022; Vohryzek et al. 2020, 2022), to investigate the mechanism of altered brain function in schizophrenia. To this aim we applied the workflow of analysis described in **Fig. 1** to a dataset including 47 patients diagnosed with schizophrenia (SCZ) and 118 healthy controls(HC) (Poldrack et al. 2016).

**Fig.1:**
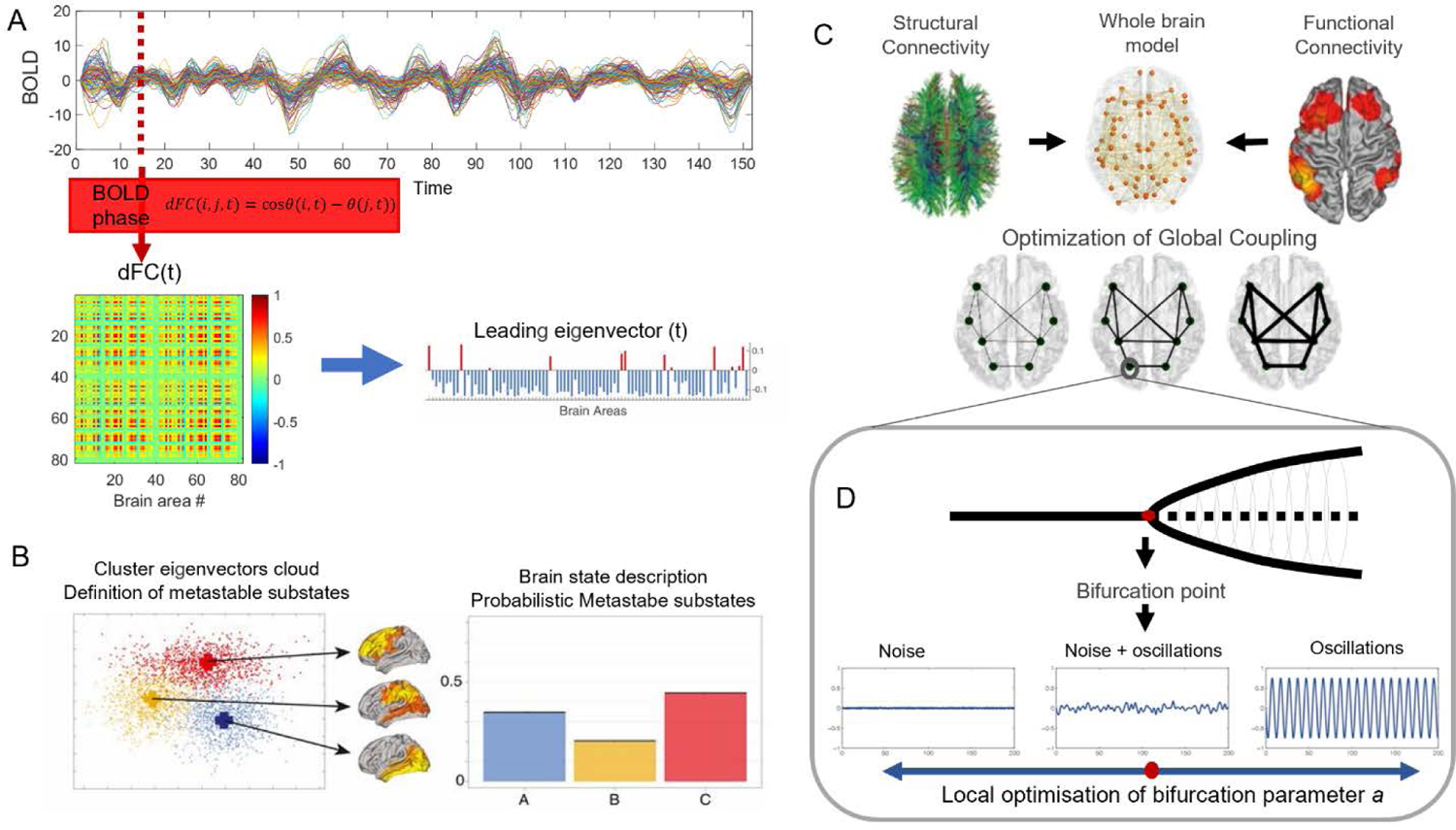
Schematic representation of the analysis workflow featuring the probabilistic metastable substate (PMS) space, whole-brain model fitting and in silico stimulations. **(A)** BOLD phases were extracted at each time point for each brain area and for each subject using the Hilbert transform. Dynamical functional connectivity (dFC) matrices were obtained as follows. For each time point the phase coherence between the BOLD signal of every pair of nodes was calculated, leading to a NxNxt matrix of instantaneous connectivity. **(B)** Leading Eigenvector Dynamic Analysis (LEiDA) was used to extract the leading eigenvector V_1_(t), capturing the most informative features of each dFC matrix. All vectors were then clustered to identify recurrent dynamic functional connectivity patterns across all subjects and conditions. Probability of occurrence of each pattern was calculated within each condition separately, defining a probabilistic metastable substate space (PMS). These probabilities were then compared between conditions. Figure adapted from Deco et al., 2019. (**C**) Structural and functional information from the empirical data was used to build a whole-brain dynamical model and fit it to reproducing the empirical PMS of the healthy and SCZ group independently. (**D**) Each brain area of the whole-brain model is systematically perturbed via in silico stimulations through two different protocols (noise and synchronization). The noise protocol shifts the local bifurcation parameter of each brain area to negative values, whereas the synchronization protocol shifts it towards positive values. Figure adapted from Deco et al., 2019.

First of all, we used LEiDA (Cabral et al. 2017) to define, identify, and compare dynamical brain substates in patients and healthy controls and to detect subtle global dynamical changes that might not be captured by global static measures. Critically, LEiDA allows us to differentiate the dynamics of the global signal from the dynamics of more focal functional pattern of activation. systems. Subsequently, we fitted a whole-brain dynamical model to both pathological and control data to reproduce those differences in-silico. Finally, we simulated the effects of regional perturbations on the healthy model to identify the most vulnerable brain areas that may be implicated in the emergence of the disease. We also applied simulated perturbations to a model of the patient data to identify areas potentially involved in compensatory process.

## 2. METHODS

### 2.1 Participants

We used the open resting-state fMRI (rsfMRI) dataset from UCLA Consortium for Neuropsychiatric Phenomics LA5c Study (Bilder et al. 2020, openneuro.org/datasets/ds000030/). We focused on data from 47 patients with schizophrenia (SCZ) and 118 healthy controls (HC). To test for disease-specific effects, we also analysed data from 49 patients with bipolar disorder (BD), and 39 with attention deficit hyperactivity disorder (ADHD), as will be explained later.

All participants gave written informed consent according to the procedures approved by the University of California Los Angeles Institutional Review Board. The dataset includes participants in the range from 21 to 50 years [33.2±9.3 ± 0.006 (mean ± SEM)]. All participants underwent extensive neuropsychological testing. Diagnoses were based on the Structured Clinical Interview for DSM-IV (SCID-I11) and on the SCID Checklist of Interviewer Behaviours and the Symptom Checklist of Interviewer Behaviours. Each group of patients (Schizophrenia, Bipolar Disorder, and ADHD) excluded anyone with one additional diagnosis among the other two, as well as with diagnosis of Substance Abuse or Dependence (not counting caffeine or nicotine), current Major Depressive Disorder, suicidality or Anxiety Disorder (Obsessive Compulsive Disorder, Panic Disorder, Generalized Anxiety Disorder, Post-Traumatic Stress Disorder). Stable medications were permitted for patients. For the control group, any previous or family history of psychiatric diagnosis, or previous psychotropic drugs exposure, were considered as exclusion criteria. Left-handed participants, as well as those who believed they might be pregnant, or had other MRI contraindications (e.g., claustrophobia, metal in body), were excluded. A detailed description of the dataset can be found in the data description by Poldrack and colleagues (Poldrack et al. 2016).

### 2.2. MRI data

#### 2.2.1 fMRI

##### Imaging acquisition

MRI data were acquired on one of two 3T Siemens Trio scanners, located at the Ahmanson-Lovelace Brain Mapping Center (Siemens version syngo MR B15) and the Staglin Center for Cognitive Neuroscience (Siemens version syngo MR B17) at UCLA. Functional MRI data were collected using a T2*-weighted echoplanar imaging (EPI) sequence with the following parameters: slice thickness=4 mm, 34 slices, TR=2 s, TE=30 ms, flip angle=90°, matrix 64 × 64, FOV=192 mm, oblique slice orientation. Additionally, MPRAGE were collected. The parameters for MPRAGE were the following: TR=1.9 s, TE=2.26 ms, FOV=250 mm, matrix=256 × 256, sagittal plane, slice thickness=1 mm, 176 slices. More information can be found at Poldrack et al., 2016.

Participants participated in two scanning sessions, one resting-state fMRI (rsfMRI) session and a second session where participants performed several behavioural tasks. For the current work, we restricted the analysis to the rsfMRI scan, which lasted a total of 304 s (T=152 time points). During the resting state participants were asked to remain still and relaxed with open eyes.

##### Pre-processing

Preprocessing of fMRI data was performed using fMRIPrep v1.1. (Esteban et al. 2019), a Nipype-based tool (Gorgolewski 2011). A non-aggressive variant of ICA-AROMA was performed on the unsmoothed outputs of fMRIPrep. Regression of mean white-matter and cerebrospinal-fluid signal were also applied. No global signal regression was applied to the data. Finally, the Desikan-Killiany atlas (Desikan et al. 2006) was used to parcellate the brain into N = 82 non-cerebellar brain areas (66 cortical areas + 16 subcortical areas). Altogether, this yielded data of size N x T (with N=82 ROIs and T=152 time points) for each participant.

#### 2.2.2 DTI

To build the structural connectivity matrix we used a connectome template available online at https://github.com/KevinAquino/HNM/find/main (see also (Deco, Kringelbach, et al. 2021)). The template was built using diffusion MRI data acquired in 289 healthy individuals using the following parameters: 2.5-mm3 voxel size; TR, 8800 ms; TE, 110 ms; FOV of 240 mm by 240 mm, and 60 directions with b = 3000 s/mm2 and seven b = 0 volumes. A single b = 0 s/mm2 volume was obtained with the reverse-phase encoding for use in distortion correction. The diffusion data were processed using MRtrix3 version 3.0 (www.mrtrix.org/) and FSL (FMRIB Software Library) version 5.0.11 (https://fsl.fmrib.ox.ac.uk/fsl/fslwiki). The diffusion images for each individual were corrected for eddy-induced current distortions, susceptibility-induced distortions, intervolume head motion, outliers in the diffusion signal, within-volume motion, and B1 field inhomogeneities. Tractography was conducted using the fiber orientation distributions (iFOD2) algorithm, as implemented in MRtrix3 (Tournier et al. 2019). To create a SC matrix, streamlines were assigned to each of the closest regions in the parcellation within a 5-mm radius of the streamline endpoints, yielding an undirected 82 × 82 connectivity matrix. A group-representative connectome was generated by selecting the 30% of edges showing the lowest coefficient of variation (based on the streamline count) across participants. Further details can be found in (Oldham et al. 2020).

### 2.3. Empirical Functional connectivity analyses

We first aimed to assess whether there were differences in FC between patients and controls at the pairwise, node, and network levels. Most frequently, FC is computed across the whole acquisition period, assuming stationarity at this time scale, which produces static functional connectivity (sFC). Such static analyses allow to keep low dimensionality but lose considerable information by overlooking fluctuations of connectivity patterns across time. Recently developed methods have been developed to capture time-varying functional couplings, defining a dynamic functional connectivity (dFC). Here, we computed both sFC and dFC and compared them as potential fingerprints of relevant alterations within our dataset. The BOLD signal was bandpass filtered the in the range 0.04-0.07 Hz to focus on the most informative frequency bands (Glerean et al. 2012).

#### 2.3.1. Static functional connectivity (sFC) and derived network metrics

For each participant, the sFC between each pair of brain areas was calculated as the Pearson correlation between their BOLD signals across the entire recording time, thus obtaining a NxN sFC matrix, where N=82 is the number of brain areas. Group averages were computed for both conditions (SCZ, HC). Pairwise differences across conditions were calculated as the group difference between the value of sFC between each pair of nodes. Global FC was calculated as the mean value of the sFC matrix across pairs of areas. In addition, we also computed each node’s strength and diversity. Strength was calculated as the sum over columns of the sFC matrix and represents the overall level of connectivity of each individual node. Diversity was calculated as the standard deviation over columns of the FC matrix to quantify the variability in each node’s connections to others. Differences in pairwise, node and global metrics across conditions were tested for significance using a Wilcoxon ranksum test with a significance level α = 0.05. Bonferroni was applied to correct for multiple comparison where necessary (N^2^ comparisons in pairwise metrics, N comparisons in node metrics).

#### 2.3.2. Dynamic functional connectivity (dFC) and LEiDA (leading eigenvector dynamics analysis)

To maintain the relevant information about the time dimension, it is possible to work with dynamic functional connectivity (dFC) instead, where the level of coupling between pair of nodes activity is measured at every time point. In our case, we defined dFC based on the phase coherence between areas, which were computed using the following procedure. We used the Hilbert transform on filtered BOLD signal to extract the phase of each brain area at each time point (**Fig. 1A**). Then, for each pair of brain areas *i* and *j* we defined the dFC as the phase coherence between them, i.e., the cosine of their phase difference at any time point *t*:

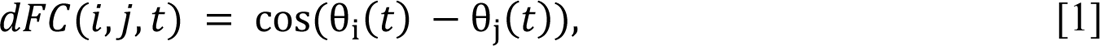

where θ_i_(t) is the instantaneous phase of the filtered BOLD signal at node i. The dFC ranges between −1 and 1. Two brain areas that are in phase (i.e., with no phase delay) will have dFC = 1, while signals in anti-phase (phase delay of π) will result in dFC=-1. In addition, orthogonal signals (i.e., with a phase delay of π⁄2) will have dFC=0. For each participant and each condition, we computed their respective dFC, which resulted in a NxNxT matrix, that allowed us to investigate the variation of connectivity pattern over time (N=82 areas, T=152 timepoints). To reduce the dimensionality of this data matrix, we extracted the leading eigenvector V_1_(t) from each dFC(t) matrix -i.e., the eigenvector with the largest eigenvalue-, thus summarising the dominant connectivity pattern at each time point in an N-dimensional vector. We thus obtained one leading eigenvector (reflecting the instantaneous FC) for each time point *t* and for each participant in each condition (SCZ and HC). Note that each element of V_1_(t) corresponds to a specific brain area and quantifies the participation of each node in the connectivity pattern. In particular, nodes can be classified as belonging to one of two communities depending on their relative sign in V_1_(t) (blue and red, see **Fig. 1B**) and the contribution to the community is represented by the magnitude of its value.

#### 2.3.3. Definition of Probabilistic Metastable Substates

Following Cabral et al. 2017, we pooled together all leading eigenvectors from all time points, participants, and conditions, and applied a k-means algorithm in order to identify recurrent brain connectivity patterns common to both conditions (**Fig. 1B**). The total number of leading eigenvectors was 25080 (T=152 time points per participant, a total of 165 participants). The k clusters defined k recurrent metastable substates in the N-dimensional space of brain connectivity substates. Each eigenvector was assigned to one of these clusters. For each cluster a centroid vector V_c_ was identified, the outer product of which represented the main connectivity pattern in that cluster. We also correlated the pattern of connectivity of each substate with the 7 functional networks as defined by Yeo’s parcellation (Yeo et al. 2011). In particular, we took the mask of the Yeo parcellation into seven non-overlapping functional networks defined in MNI152 space1 and the mask of the DK parcellation in the same MNI152 space, and calculated, for each of the 82 brain areas, the proportion of voxels assigned to each of the seven functional networks, obtaining seven (1×82) vectors representing the intrinsic functional networks in DK space. Subsequently, we computed Pearson’s correlation between these seven networks and the centroids obtained from our clustering analysis (setting all negative values of the centroids’ vectors to zero, to consider only the areas whose BOLD phase is shifted from the main orientation).

The analysis was run repeatedly with cluster number k varying from 2 to 15. After running the clustering for all the values of k we focused our analysis on k=3, which resulted in the minimal number of substates to reflect differences between conditions SCZ and HC. In addition, k=3 has been shown in previous research to offer a good characterisation of brain dynamics (Deco et al. 2019; Escrichs et al. 2022; Vohryzek et al. 2022). It allows to reach a good balance between informativity and synthesis, important to increase meaningful interpretability of the results by focusing on the most relevant dynamics. Keeping the dimensionality low also allows us to replicate more easily the empirical results with a phenomenological in silico model, and proceed with the investigation of critical areas responsible for such dynamics.

We then aimed to characterise the recurrent behaviour of the three identified common metastable substates by investigating the way in which they alternate over time in patients and controls. To do so, we pooled together all leading eigenvectors from all time points and participants for each condition separately. For each metastable substate (m=1,2,3), we estimated its probability of occurrence P(m), as the number of eigenvectors assigned to cluster m divided by the total number of eigenvectors. This defined a “probabilistic metastable substates (PMS) space” for each condition, consisting of the overall distribution pattern of the probabilities of occurrences of the substates. Note that, by definition, when this procedure is applied to data that have not undergone global signal regression, as in the current case, the most frequently recurring pattern of connectivity identified is represented by the global state, while the others represent specific functionally oriented pattern of activation (Vohryzek et al. 2020). Differences between the probability of occurrence (PO) in SCZ and HC were tested for significance by using a permutation test to build a surrogate distribution of POs for each substate m=1,2,3 under the null hypothesis that there is no difference between conditions. For each substate m, our test statistic was defined as the difference in PO across conditions: Δ(m)=P(m)-Q(m), where P(m) and Q(m) stand for the PO of state m in the control and schizophrenia groups, respectively. Then, we generated 1,000 independent samples of the differences Δ(m) using the following procedure. Eigenvectors from all participants, time points and conditions were pooled together. Condition labels (schizophrenia, control) were randomly shuffled across all eigenvectors, while keeping cluster labels (1,2,3) unchanged. We then compared each difference Δ(m) (m=1,2,3) with its corresponding surrogate null distribution and computed the p-value of the hypothesis test. The significance level was set to α = 0.05, p-values were corrected for multiple comparisons (k=3 comparisons).

#### 2.3.4. Time-average dFC and derived network metrics

To understand whether differences in the recurrence of states between conditions resulted in altered time-persistent connectivity, we also computed the time-averaged dFC for each participant. Similar to the sFC, the time-averaged dFC is an NxN matrix which quantifies the degree of coupling across time. However, while the former relies on signal co-fluctuations captured by Pearson correlation, the latter reflects phase synchronisation sustained across time. From the time-averaged dFC, individual node strength, node diversity and global FC were computed following the procedure described in Section 2.3.1. We also tested for significance in pairwise, node and global differences using Wilcoxon ranksum tests, with a significance level α = 0.05. Bonferroni was applied to correct for multiple comparison where necessary. Metastability, a measure of the variability of global synchronization, was computed as the standard deviation of the Kuramoto order parameter, which measure the level of phase coupling between signals across time.

### 2.4. Whole-brain network model

Here, we used a whole-brain generative model capable of replicating the dynamics of the metastable substates revealed by the previous analysis to simulate the response of the system to external perturbations. The whole-brain model was fitted using the structural and functional connectivity data derived from neuroimaging, as will be described later.

Within this model, each of the *N* = 82 nodes are modelled as a Stuart-Landau oscillator, which in turn are coupled through the empirical structural connectivity. Specifically, the local dynamics of each brain area *i* is described using the normal form of a supercritical Hopf bifurcation with bifurcation parameter *a*_*i*_ (**Fig. 1C**). According to this model, a bifurcation occurs at *a*_*i*_ = 0, so that for *a*_*i*_ < 0 the node displays noisy low-amplitude asynchronous activity, and for *a*_*i*_ > 0 a limit cycle exists, and the node displays oscillations with intrinsic frequency *f*_*i*_ = ω_i_/2π (*i* = 1, …, *N*). In the current work, we set each node’s intrinsic frequency within the range 0.04-0.07 Hz, which has been repeatedly shown to be the most functionally relevant (Glerean et al. 2012) The frequency ω_i_ was estimated from the empirical data, as given by the averaged peak frequency of the narrowband BOLD signals of each brain region *i* = 1, …, *N*.

The coupling between the different nodes is built to reflect the structural connectivity derived from diffusion MRI and the strength of these connections is regulated by a global coupling parameter G. The whole-brain dynamics can therefore be described by a set of coupled differential equations, where for each node *i* = 1, …, *N* we have:

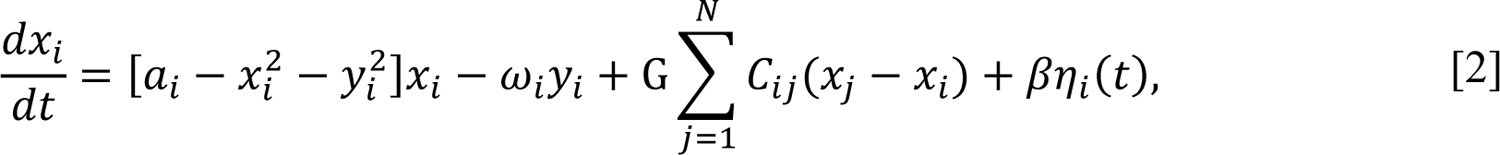

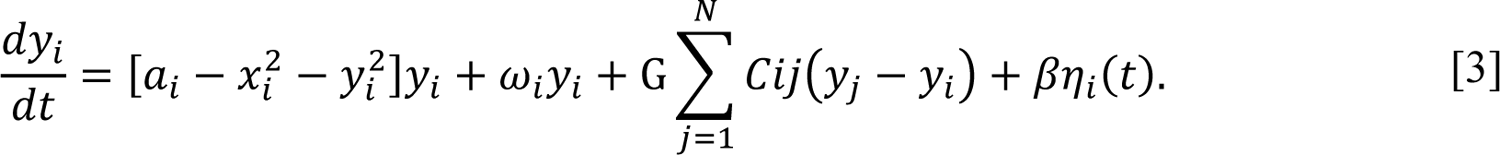

where the variable *x*_*i*_ models the BOLD signal of area *j, η*i**(t) is additive Gaussian noise with standard deviation η and C_*i*j_ denotes the density of fibres between areas *i* and j as given by the DTI. The noise η was fixed to be equal to 0.02 and the mean SC matrix *C* = (*C_*i*j_*) was normalised by a factor of 0.2 in order to be in the same range of parameters explored in Deco et al. (2017). Since in that same work the optimal working point of the model for resting state brain activity has been shown to be at the edge of the Hopf bifurcation, we set *a*_*i*_ = 0 in each node *i* and we modified the parameter G allowing it to vary from 0 to 0.50 in steps of 005, to find the best fit in each condition separately.

To evaluate the goodness of fit of the model at each value of G we used two complementary measures. First, we compared empirical and simulated FCD matrices by quantifying the distance between the cumulative distribution functions of the two samples by computing the Kolmogorov-Smirnov (KS) statistics. For a single participant session where T time points were collected, the corresponding phase-coherence based FCD matrix is defined as a TxT symmetric matrix whose (t1, t2) entry is defined by the cosine similarity between the upper triangular parts of the 2 matrices dFC(t1) and dFC(t2) (previously defined; see above).

Additionally, we compared empirical and simulated probabilities of occurrence P(m), where m=1,2,3, as previously defined in the PMS. Specifically, for each value of G we simulated BOLD signals using equations 2 and 3 and the parameters specified above. Once BOLD signals were generated, LEiDA analysis was applied to the simulated data, and the empirical centroids were used to cluster the pattern of activity extract the probability of occurrence P(m) for each substate m=1,2,3. To compare the probabilities of occurrence for the empirical and simulated data, we used a symmetrised Kullback– Leibler divergence (aka. KL distance):

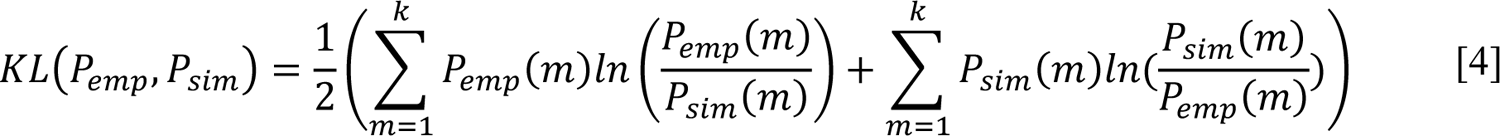

We ran 50 simulations for each value of G and computed the mean KL and KS distance across iterations. We considered the value of G where mean KL distance was minimised as the optimal working point of the model. For that value of G, we computed the averaged simulated PMS across simulations. More information about the model and the fitting procedures can be found in (Deco et al. 2019).

### 2.5. Perturbation

Once the whole-brain model had been optimised for each condition separately, we proceeded to systematically alter the activity of each individual node to highlight critical areas that might induce a state transition from one condition to the other as measured by the probability of occurrence of each metastable substate.

First, we studied perturbations applied on the healthy condition (source condition) and quantified changes in the PO distribution. Perturbations were applied bilaterally for each of the 41 pairs of nodes. For each pair of nodes, we gradually changed their local bifurcation parameter a, while keeping every other pair of nodes at the original value a=0. The perturbation was applied for each pair of nodes in both directions, i.e., towards increased noise (a<0) and towards increased synchronisation (a>0). The values of a ranged from 0 to −0.3 in steps of −0.015 for the noise paradigm and from 0 to 0.2 in steps of 0.01 for the synchronisation paradigm, according to previous research (Deco et al. 2019). For each combination of parameters (perturbed area, intensity of perturbation), we simulated BOLD signals using the model described above and evaluated the effectiveness of the transition by computing the KL distance between the PMS of the perturbed source state (HC) and the target state (SCZ). We repeated this procedure 50 times and obtained the mean and standard deviation of the KL distance across simulations for each combination of parameters. We then built a baseline KL distance distribution by pooling all KL distance values across simulations and areas for a=0, i.e. distribution of KL distances between non-perturbed simulated HC and empirical SCZ across simulations, and used this distribution to z-score all values in the matrix. We then identified as critical the areas where perturbation induced the greatest reduction in KL distance.

Additionally, we investigated the alterations of global static measures of functional connectivity as a consequence of local perturbation of sensitive areas. To this aim we compared levels of pairwise and global functional connectivity, strength, diversity and metastability corresponding to data generated by the model at different values of bifurcation parameter. Differences were tested for significance using a Wilcoxon ranksum test across simulations with a significance level α = 0.05. Bonferroni was applied to correct for multiple comparison where necessary (NxN comparisons in pairwise metrics, N comparisons in node metrics).

Finally, we repeated the whole procedure described above inverting the direction of the transition. We therefore used the PMS simulated through the model reproducing the SCZ condition as source state and the empirical PMS of the HC condition as the target state.

## 3. RESULTS

### 3.1. Defining probabilistic metastable substates

We first analysed dynamical properties of the empirical data and looked for relevant differences between conditions (patients with schizophrenia, SCZ; healthy controls, HC) through leading eigenvector dynamics analysis (LEiDA), as shown in **Figs. 1A** and **1B**. This methodology allowed us to identify the dominant pattern of connectivity, define functional communities at each time point, and analyse the dynamics through which these patterns alternate over time (Cabral et al. 2017). We then used k-means clustering to define the main recurrent substates, each of which represented a connectivity pattern (defined by its centroid) with an associated probability of occurrence (PO). The distribution of probabilities of the substates determined the so called “probabilistic metastable substates (PMS) space”, that can be used as a definition of each specific state condition (Deco et al. 2019) The procedure was applied to concatenated data of participants including both healthy controls and patients with schizophrenia, in order to identify common substates and allow for comparison.

We first ran the analysis for different numbers of clusters (k=2, …,14); k=3 was found to be the minimal number of clusters to capture differences in PO between cases and controls (see methods). We therefore restricted the study to the analysis of the dynamical properties of the principal three main substates. **Fig. 2A** depicts the centroid vector of each of the three main substates (V_c_), each characterising a functional connectivity organisation across brain areas, as represented in the corresponding FC matrix. To facilitate interpretation, we also projected the results of the analysis in the brain (**Fig. 2B**) and computed the functional connectivity matrix corresponding to each substate (**Fig 2C**). Finally, **Fig. 2D** represents the correlation of the connectivity pattern of each substate with the 7 functional networks as defined by Yeo’s parcellation.

**Fig.2:**
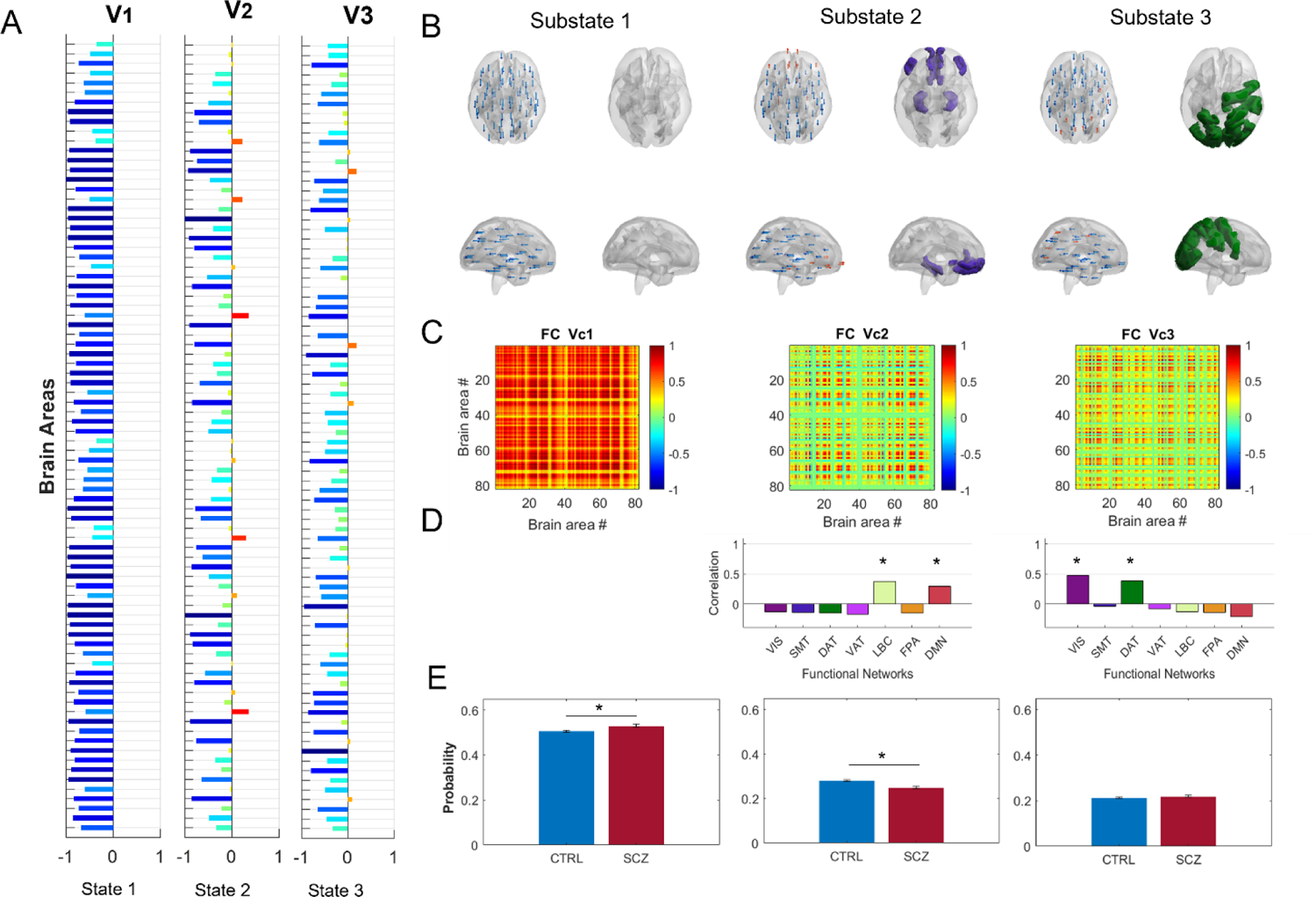
Empirical PMS and group comparison. Probabilistic metastable states (PMS) were identified through LEiDA analysis, by applying a k-means clustering algorithm to all leading eigenvectors from all subjects and conditions together. The procedure was repeated for k = 2 to 15 clusters. K=3 resulted to be the minimal number of substates to reflect differences between conditions SCZ and HC. **(A)** Leading eigenvectors V_1_(n) representing the pattern of connectivity of the centroids of each metastable state. Two network communities (scale of blues and scale of reds) of nodes(n) are identified depending on the respective sign in V_1_(n) and the contribution to the community is represented by the magnitude of his value. **(B)** Areas that positively coactivate in each substate are plotted in the brain to facilitate visualisation, and **(C)** the functional connectivity matrix of each respective centroid is shown. **(D)** For each substate we explored correlation with Yeo networks. The first substate is characterised by global synchronisation across all nodes. Hence, it has no correlation with any specific functional network and represents the global signal. The second substate positively correlates with the default mode network and the cortical limbic network, and may reflect internally oriented states. The third substate and correlates with visual network and the externally oriented attention network, and may reflect externally oriented states. **(E)** Probability of occurrence of each substate was computed and compared across conditions. The bottom panel shows the empirical PMS of each condition (in red the group of patients with schizophrenia and in blue the healthy group). The probability of occurrence in the first substate was significantly higher (p<0.01) in the SCZ group (0.531 ± 0.006; mean ± SEM) than in the HC group (0.506 ± 0.004). By contrast, the probability of occurrence in the second substate was significantly higher (p<0.01) in the control group (0.288 ± 0.003) than in the SCZ group (0.250 ± 0.004). No difference between condition was found in regard to the PO of the third substate.

The most frequently recurring substate is characterised by global synchronisation of the whole brain, as shown by all areas being partitioned into the same community according to the sign of the leading eigenvector, and by the presence of positive value of correlation across the whole FC matrix. This substate represents timepoints wit elevated amplitude of global signal, defined as a widespread level of synchronisation across all voxels. The other two substates, in contrast, represent the emergence of specific functional connectivity patterns. In particular, the second substate is characterised by the organisation of associative areas, mainly prefrontal and parahippocampal, into a functional community whose dynamics oppose to the rest of the brain. This pattern of activation significantly correlates with default mode (r=0.42, p<0.05) and limbic networks (r=0.32, p<0.05) and can be summarised as a top-down internal control substate. In contrast, the third substate is characterised by the emergence of a functional community formed by areas involved in visual processes and sensorimotor integration (externally driven attention), and significantly correlates with visual (r=0.52, p<0.05) and dorsolateral attention (r=0.38, p<0.05) networks.

### 3.2. Differences in the PMS space between conditions

For each of these substates we computed the probability of occurrence (PO) over time and analysed relative differences between conditions (**Fig. 2E**). We found that the PO of the first metastable substate was significantly higher in the SCZ group (0.531 ± 0.006; mean ± SEM) than in the HC group (0.506 ± 0.004). Significance was tested by means of a permutation test shuffling condition labels across all time points and participants (p<0.01, FDR-corrected for k=3 comparisons, Cohen’s D effect size D = 5). These results indicate an augmented dominance of the global signal over other dynamics in the pathological group, in line with previous research showing an increase in power spectrum and variance of global signal in this condition (Yang et al. 2014, 2017), confirming that it is a relevant characteristic of this pathology.

By contrast, the second metastable substate’s probability was significantly lower in the SCZ group (0.249 ± 0.004) when compared with the HC group (0.281 ± 0.003, p<0.01, FDR-corrected for k=3 comparisons, Cohen’s D effect size D = 11). This suggests a reduction of the rate at which synchronisation within a fronto-temporal and subcortical associative network dominates the overall brain dynamics in patients as compared to the healthy controls. No difference was found in the PO of the third substate between conditions (0.212 ± 0.005 in SCZ vs. 0.210 ± 0.003 in HC, p=0.2, FDR-corrected for k=3 comparisons).

Given the limited knowledge on how global signal varies across conditions and the debate on the use of GSR, we wanted to test whether the differences that we identified were specific to schizophrenia, rather than being generically associated with psychiatric disorders. To test this hypothesis, we repeated the same analysis with a cohort of 49 participants with bipolar disorder (BD) and a cohort of 39 participants with attention-deficit/hyperactive disorder (ADHD) from the same dataset, and we compared them with the same healthy control (HC) group used for the previous analyses. In neither case did we find a significant difference between the PO of the first substate (reflecting the global signal) in the pathological group compared to HC, although we found significant differences in the other two substates (supplementary **Fig. S1**). This confirms previous results indicating the specificity of the changes in GS associated with schizophrenia, suggesting it is a diagnostically specific finding (Yang et al. 2014).

Complementary to the main analysis, we investigated whether these changes in dynamical connectivity could be explained by or reflected in static or time-persistent alterations in the network connectivity. To this aim we looked for differences in network metrics derived from two complementary FC measures: the sFC, based on Pearson correlations, and the time-average dFC, based on the consistency of phase synchronisation across time. For each FC measure, we investigated differences in pairwise FC, node FC strength, node FC diversity and global FC between SCZ and HC. No differences were found between these measures across patients and controls (supplementary **Fig. S2**), suggesting that alterations in the system might be better captured by measures targeting the temporal structure of a system that is intrinsically dynamic.

### 3.3. Reproducing the empirical differences with a whole-brain network model

The next step of the work consisted in fitting a whole-brain model to the PMS of each condition. To this end, we used a generative model as previously described in others work (Deco et al. 2017, 2019) to simulate the activity of *N* = 82 nodes, linking structural and functional information derived from the empirical data and adjusting the free parameters to achieve the best fit. The local dynamics of each node *i* were described with the normal form of a Hopf bifurcation with an intrinsic frequency ω_*i*_ derived from the empirical data. Based on previous research, each node was set to be working at the verge of a Hopf bifurcation, as defined by the local bifurcation parameter *a*_*i*_ = 0 (*i* = 1, …, *N*). This results in a noisy-oscillatory behaviour. Furthermore, the model allows the activity of each node to evolve depending on the dynamics of the others, according to SC constraints derived from diffusion MRI. A more detailed description of the model can be found in the method section.

To search for the optimal fit, we adjusted the global level of coupling by exploring different values of the free parameter G, which controls for the conductivity of the fibres given by the SC by scaling the strength of the interarea connectivity (**Fig. 3A**).

**Fig.3:**
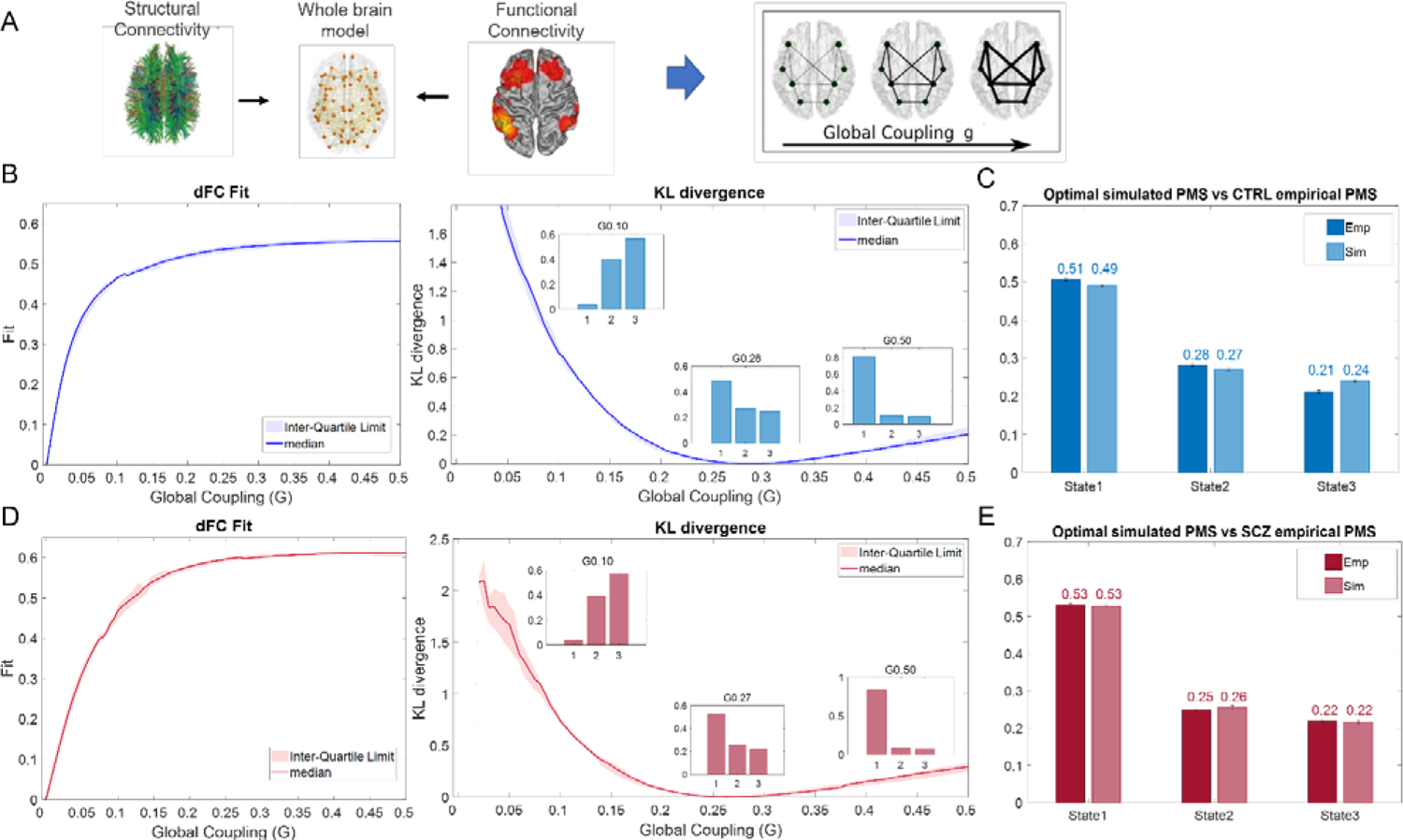
Whole-brain and PMS model fitting. **(A)** Structural and functional information from the empirical data were used to build two whole-brain dynamical model. For each condition simulated BOLD signal was generated through the model and the simulated PMS was computed. By adjusting the global parameter G, an optimal fit for the PMS was obtained for the HC and SCZ groups, separately. **(B)** The left panel shows the level of FCD fit of the model for each value of global coupling G for the healthy condition (shaded area represents the inter-quartile range across N=50 simulations). The right panel shows the values of Kullback–Leibler (KL) distance between the empirical and the simulated PMS). Examples of simulated PMS are shown to appreciate the corresponding change in the PO of the three substates at different values of coupling. **(C)** The panel shows the resulting optimal simulated PMS (light blue), which best fits their empirical counterpart for the healthy group (dark blue) **(D)** The left panel shows the level of FCD fit of the model for each value of global coupling G for the SCZ condition, while the right panel show the values of KL distance between the empirical and the simulated PMS. **(E)** The panel shows the resulting optimal simulated PMS (light red), remarkably similar to their empirical version for the schizophrenic group (dark red). Error bars represent SEM across subjects for the empirical PMS and SEM across simulations in the simulated PMS.

The optimization was done independently for each group. For each value of G, BOLD signals were simulated using the whole-brain model. Simulated dynamical functional connectivity matrices were computed and LEiDA analysis was applied as previously described to investigate the dynamics of the metastable substates in the simulated data. Through this procedure we were able to accurately reproduce in silico the empirical dynamics of the HC condition. **Fig. 3B** shows the level of fit of the model as a function of the global parameter G measured both with the dFC fit and the KL distance (Kullback–Leibler divergence). In particular, as the KL distance quantifies the difference between the empirical and the simulated PMS, a low value of this measure represents a good level of accuracy of the model in replicating the empirical data. In our case, the optimal fit was found at a value of G = 0.287 ± 0.012, with KL = 0.003 ± 0.001 (mean ± SEM for N=50 simulations). The ability of the model to reproduce the empirical dynamics at the optimal working point can be also appreciated in **Fig. 3C**, which shows the PMS simulated by the model (P(1)=0.489±0.003, P(2)=0.269±0.002, P(3)=0.240±0.002, mean ± SEM for N=50 simulations). These values replicate well the corresponding empirical ones (P(1)=0.506 ± 0.004, P(2)=0.281 ± 0.003, P(3)=0.212 ± 0.003, mean ± SEM for N=127 participants).

We were also able to accurately reproduce in silico the empirical dynamics of the SCZ group and to accurately replicate the differences between the two conditions as shown in **Fig. 3D**. The optimal fit was found at a value of G=0.27 ± 0.02. At this value of coupling in the model was able to accurately replicate the empirical data with KL=0.002 ± 0.002 (mean ± SEM for N=50 simulations). The corresponding probability of occurrence of the each metastable substate were P(1)=0.527±0.004, P(2)=0.255±0.003, P(3)=0.217±0.002 (mean ± SEM for N=50 simulations), which accurately replicated the empirical ones (P(1)=0.531 ± 0.006, P(2)=0.249 ± 0.004, P(3)=0.219 ± 0.005, mean ± SEM for N=47 participants).

### 3.5. Perturbing the model to identify vulnerable areas potentially underlying the emergence of the pathology

Next, we investigated whether it was possible to induce a change in brain state (schizophrenia and healthy)–as defined by the PMS– by applying a perturbation to the model. Specifically, we inquired whether we could simulate the pathological-like behaviour of the SCZ condition by changing only the bifurcation parameters of the model optimised to reproduce the HC empirical data (**Fig 4**).

In addition, we wanted to explore how the effect of the perturbation changed as a function of the targeted ROI. Specifically, we aimed to identify nodes that could induce a global change at the whole brain level sufficient to determine a significant shift in the PMS space in a way that would resemble the pathological one.

To this aim, we systematically perturbed each bilateral pair of nodes (N=41) in the simulated model of the healthy control condition by changing the value of the local bifurcation parameter (*a*), progressively moving it toward more negative (0 to −0.3) or positive values (0 to 0.2), with steps of 0.01. In a second level analysis, we repeated the perturbation decreasing the inter-step distance to 0.0025 to have better resolution in the range 0 to 0.05, as it resulted to be of particular interest as described below.

Decreasing the value of the local parameter did not lead in any case to a change in the PMS that resembled more the pathological condition, and in certain cases it increased the difference even more. Fig. 4A shows the change in KL distance induced by the perturbation (expressed a z-score with respect to the non-perturbed starting point (see methods). We can observe that as soon as a minimal perturbation is applied to any brain area in the direction of more negative values of the parameter *a*, the delta KL starts to increase, reflecting a worsening of the fit with the target state (SCZ). Additionally increasing the intensity of the perturbation induced a progressive further worsening of the fit. A few areas were less affected by the perturbation and induced only a moderate increase in KL distance, but never led to a decrease.

**Fig.4:**
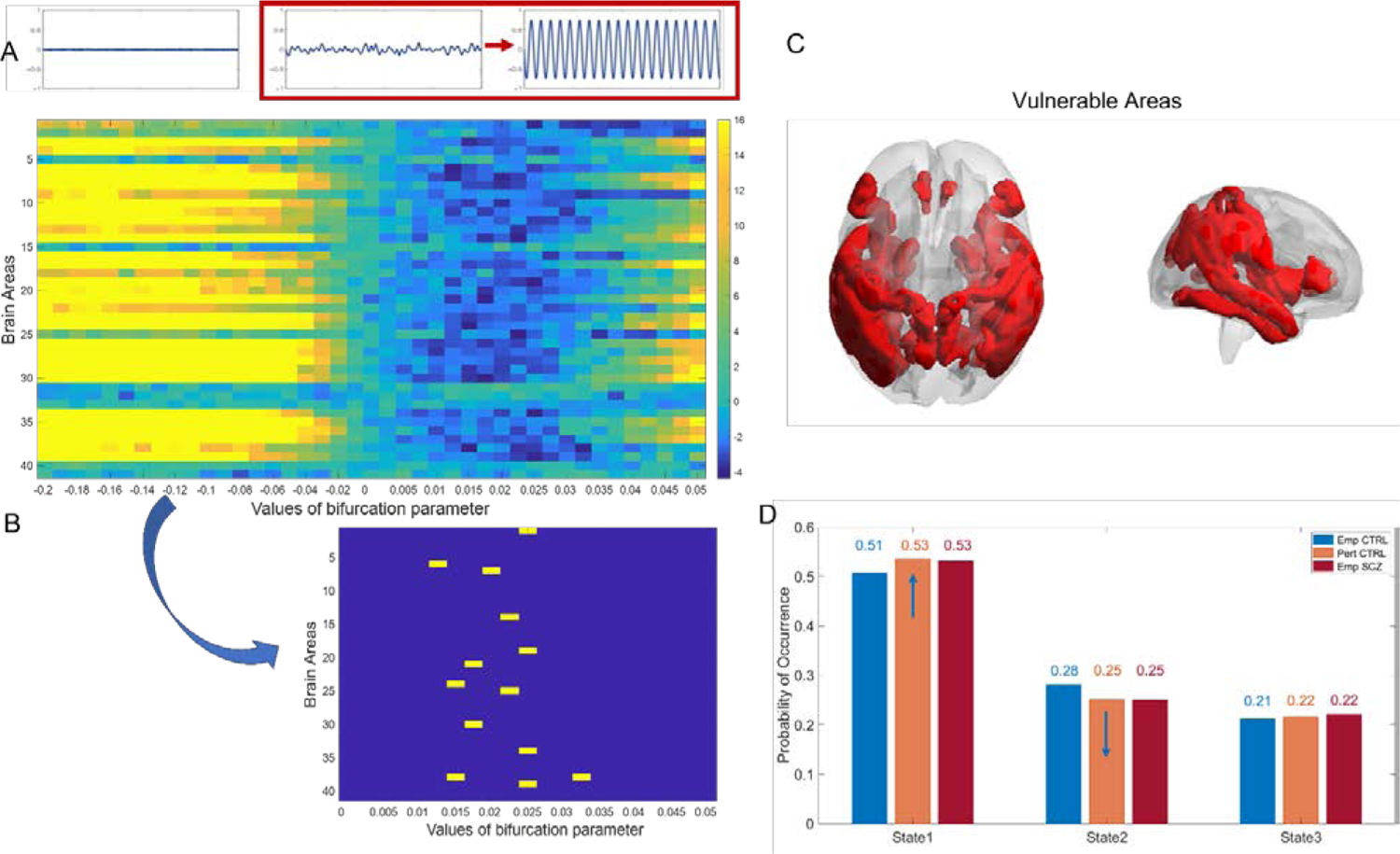
Perturbation protocols from the healthy to the pathological condition. **(A)** Each pair of bilateral nodes in the whole-brain model of the healthy condition was perturbed individually at the optimal working point by shifting the local bifurcation parameter to negative and positive values. The optimal perturbation is that which achieves that the first brain state decreases, the second increases, and the third brain state remains similar. The matrix shows the KL-distance between the empirical PMS of the schizophrenia group (target) and the perturbed model of the healthy condition (source) after applying the perturbation protocols for different intensity (from softer to stronger) in both directions in each brain area. Values are expressed as a z-score with respect to a baseline non-perturbed (a=0) distribution, built by pooling all KL-distance values across nodes and simulations. We can notice how the fit improve shifting the bifurcation parameter a towards more positive values, and therefore increasing the level of internal synchronisation and oscillatory behaviour. **(B)** The most vulnerable areas were identified by thresholding the 1.5 percentile of KL distance and selecting areas where the perturbation was more effective in inducing a global shift toward the pathological condition. **(C)** The vulnerable areas were rendered onto the cortex to facilitate visualisation and represent brain areas potentially involved in the emergence of the disease. **(D)** PMS comparison, showing the transitions from the model of the healthy control group (blue) to the empirical PMS of the SCZ group (red), as a result of the perturbation of inferior temporal cortex (orange).

On the contrary, bilaterally perturbing brain regions by shifting the corresponding local bifurcation parameter *a* toward positive values induced a global shift in the simulated PMS state. In fact, it progressively reduced the KL distance with the empirical data of the pathological condition until we could observe a change of state, by increasing the PO of the global state and decreasing the PO of the second substate so that the simulated PMS resembled the PMS of target state (SCZ) more than the one corresponding to the source state (HC), as can be seen in **Fig. 4D**. In particular we observe in **Fig 4A** that the ΔKL decreases, reflecting an improvement of the fit with the target state (SCZ), when a perturbation is applied in the range of values between *a* = 0.005 and *a* = 0.035. For more intense values of perturbation, the fit remained good only for few areas while it worsens for the others.

We identified the most vulnerable areas defined as the ones whose perturbation induced the most significant shift. In particular, we selected the 1,5% areas most responsive to perturbation. This resulted in selecting those areas where we could detect an improvement in KL distance equal or superior to −3.7503 standard deviations away from the non-perturbed starting point (**Fig. 4B**, Suppl. **Table1**). To facilitate interpretation, we plotted such areas in the brain (**Fig. 4C**). The areas that induced the most significant shift toward the pathological condition are primary sensory areas like postcentral cortex, as well as associative areas, involved in the processing and integration of external stimuli, such as the insula, inferior parietal, fusiform cortex and middle temporal cortexes or in memory like the hippocampus. We show an example of effective perturbation in the inferior temporal cortex, which causes a particularly large reduction in KL (with target) (ΔKL=-3.79, KL=0.0012+-0.0003). We illustrate how, by changing the bifurcation value in this area, the global PMS state changes significantly and finally resembles the pathological data (**Fig. 4D**). In addition, we find that the shift towards the pathological condition induced by the perturbation increases global FC (p<0.01) (data not shown).

### 3.6. Perturbing the model to identify sensitive areas potentially involved in compensatory mechanisms

Finally, we repeated the same procedure in the opposite direction to identify critical areas potentially involved in compensatory mechanisms that could induce a shift in the model toward a healthy-like state. We systematically perturbed each bilateral pair of nodes (N=41) in the simulated model of schizophrenia and evaluated how the perturbations influenced the KL distance with the empirical data of healthy participants.

To induce a shift toward a healthy-like functioning, the bifurcation parameter had to be tuned towards negative values, progressively decreasing oscillatory behaviour. We saw that bilaterally perturbing brain regions by decreasing their values of *a*, significantly altered the global dynamics. Specifically, it induced a progressive decrease of the dominance rate of the global state and an increase of the probability of occurrence of the second substate, representing prefrontal-control, until it was possible to observe a simulated PMS that highly resembled the empirical one for the healthy control group better than patients’ data (**Fig 5**).

**Fig.5:.**
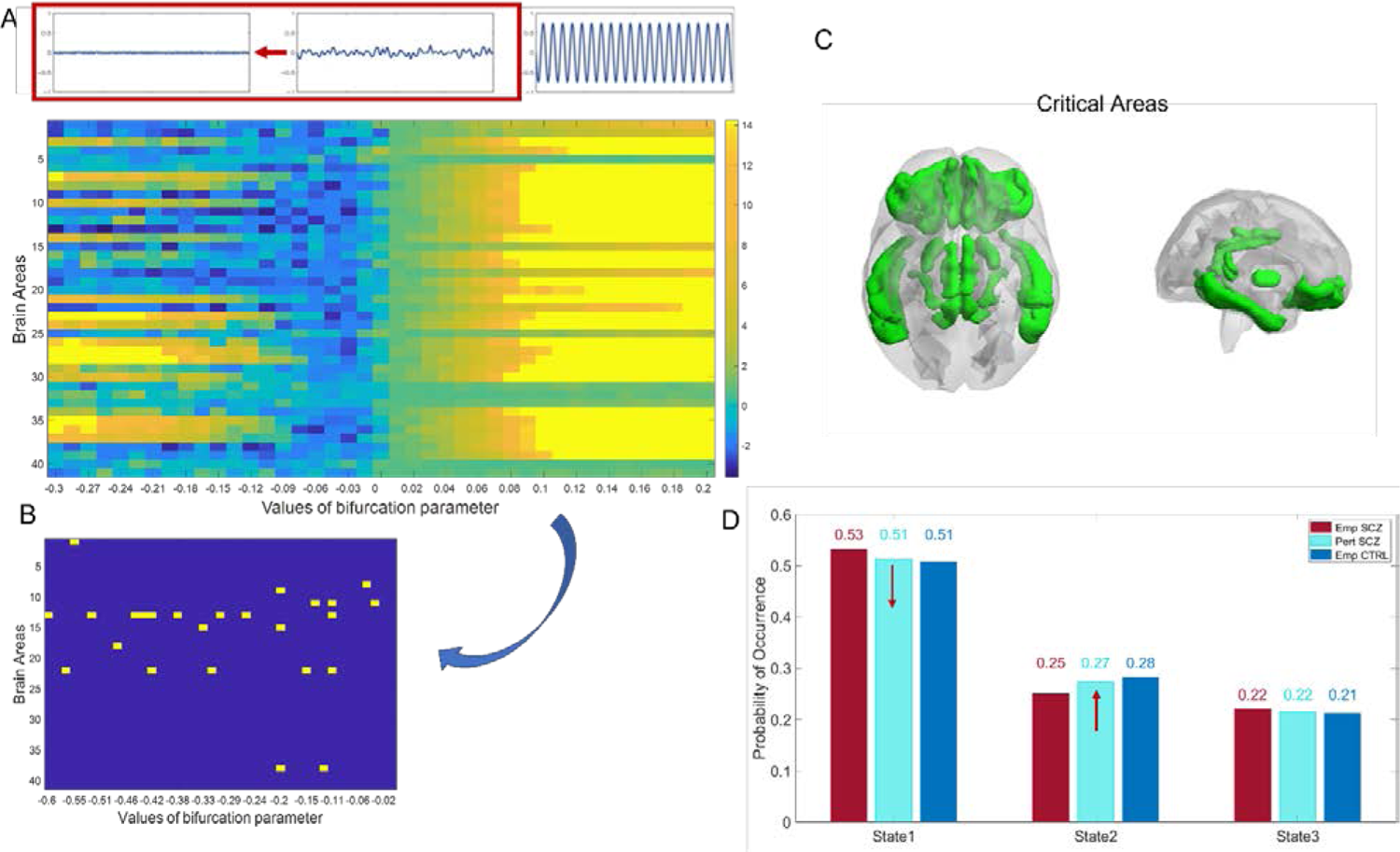
Perturbation protocols from the pathological to the healthy condition. **(A)** Each pair of bilateral nodes in the whole-brain model of the pathological condition was perturbed individually at the optimal working point by shifting the local bifurcation parameter to negative and positive values. The optimal perturbation is that which achieves that the first brain state increases, the second decreases, and the third brain state remains similar. The matrix shows the KL-distance value between the PMS of the healthy group and the perturbed model of the schizophrenia condition after applying the perturbation protocols for different intensity (from softer to stronger) in both directions in each brain area. We can notice how the fit improve shifting the bifurcation parameter a towards more negative values, and therefore reducing the level of internal synchronisation and oscillatory behaviour. **(B)** The most vulnerable areas were identified by thresholding the 1.5 percentile of KL distance and selecting areas where the perturbation was more effective in inducing a global shift toward the healthy condition. **(C)** The vulnerable areas were rendered onto the cortex to facilitate visualisation and represent brain areas potentially involved in the emergence of the disease. **(D)** PMS comparison, showing the transitions from the model of the schizophrenic group (red) to the empirical PMS of the healthy group (dark blue), as a result of the perturbation of medial orbitofrontal cortex (turquoise).

In particular we can observe in **Fig 5A** that when perturbation is applied in the direction of more negative values of the parameter *a*, the delta KL starts to decrease, reflecting an improvement of the fit with the target state (HC). Individual perturbation of the majority of areas induced a decrease in KL distance starting from *a* = −0.03 until *a* = −0.09. For more intense values of perturbation, the fit kept increasing for most areas while it worsens back for the others.

As before, we identified the most critical areas defined as the ones whose perturbation induced the most significant shift. We applied the same threshold used in the previous analysis and selected the 1,5% areas most responsive to perturbation. That resulted in selecting those areas where we could detect an improvement in KL distance equal or superior to −2.9828 standard deviation away from the non-perturbed starting point (**Fig 5B**, Suppl. **Table2**). Again, we plotted such areas in the brain to facilitate interpretation (**Fig 5C**). The nodes that induced shifts in the model toward the healthy state are mainly prefrontal areas such as medial and lateral orbitofrontal cortex and high associative areas involved in internal processes like memory or language processing, such as posterior-cingulate, isthmus cingulate, IFG (pars orbitalis), parahippocampal. We show an example of effective perturbation, illustrating how, by changing the bifurcation value in the medial orbitofrontal cortex, which causes a particularly large reduction in KL (KL= 0.0016+-0.0004, ΔKL = −3.47), the global PMS state changes significantly and finally get to resemble the healthy one (**Fig 5D**). Accordingly, increasing the value of the local parameter did not lead in any case to a change in the PMS that resembled the healthy condition, and in most cases, it actually worsened the fit. As we can observe in **Fig 5A**, as soon as a minimal perturbation is applied to any brain area in the direction of more positive values of the parameter *a*, the delta KL starts to increase, reflecting a worsening of the fit with the target state (HC). A few areas were less affected by the perturbation and induced only a moderate increase in KL distance. Finally, the superior temporal sulcus, the IFG (pars triangularis) and the pallidum were associated with shifts in both directions, from the healthy to the pathological state and vice versa.

### 3.7. An example of effective perturbation: the medial orbitofrontal cortex

To better understand the mechanisms underlying the state change induced by the decrease of the value of the bifurcation parameter, we deepened the analysis on the effects of the perturbation of the medial orbitofrontal cortex. This area was particularly responsive to perturbation and has been shown to be relevant in schizophrenia (Demjaha et al. 2022; Waltz and Gold 2007), with increased FC between the OFC and subcortical areas being identified in psychosis patients and their first degree relatives (Fornito et al. 2013). We therefore analysed how several static global measures were affected as a result of the perturbation of this area. We found that the perturbation significantly decreased (*a*=0 vs *a*=-0.1 and *a*=-0.42; p=0.03, p<0.01) the strength of the target area (Supplementary **Fig. S3A**). Moreover, it induces a significant (*a*=0 vs *a*=-0.1 and *a*=-0.42; p<0.01, p<0.01), albeit less consistent, decrease of the global FC of the whole brain (Supplementary **Fig. S3B**). This would indicate that the increased FC of medial orbitofrontal cortex is a relevant feature for schizophrenia, and that decreasing it can potentially contribute to restore global healthy levels of connectivity.

Metastability levels were not affected by the perturbation (Supplementary **Fig. S3C**). At the level of pairwise functional connectivity, perturbation of medial orbitofrontal cortex induces a more complex combination of pairwise FC alterations. The connectivity in most of the pairs of brain areas did not change with perturbation, but in a minority of pairs we detected a significant change (p<0,05). None of these values survived multiple comparison corrections. Within the affected areas, most decreased FC as a result of the perturbation, but in some pair of nodes, mainly involved in sensory processing and memory, FC increased. This suggests a non-trivial effect of perturbation on dynamical alterations (Supplementary 3D). This result also shows that the global change was driven by a minority of circuits where FC was significantly affected by the perturbation.

## 4. DISCUSSION

In the current study we investigated dynamical alterations of the functional organisation of the brain in schizophrenia. In particular, we applied dimensionality reduction and a clustering approach on a measure of dynamical functional connectivity to accurately separate the alterations related to globally coherent activity from alterations in specific pattern of activation. We then used *in silico* perturbation of a whole-brain model to identify areas involved in the emergence of the disease and in potential compensatory mechanisms. Our approach allowed us to characterize the pathological and healthy conditions (HC, SCZ) identifying the corresponding probabilistic metastable substates (PMS) space, and to reproduce the crucial alterations in silico. We could separately study and interpret differences within global coherence state from other relevant differences within the rest of the signal. Particularly, we could show that the group of patients with schizophrenia was characterised by both alterations in global signal and in specific functional networks, confirming past findings (Li et al. 2019; Yang et al. 2014). Our perturbational approach additionally allowed us to identify critical areas crucially involved in determining relevant healthy and pathological brain dynamics.

### Schizophrenia is characterised by alterations in functional dynamics of the global signal

Several studies have found alterations in GS, although the results are not always consistent. For instance, Farinha et al. described decreased GS and increased recruitment of functional networks in schizophrenia (Farinha et al. 2022). On the other hand, increased cortical power and variance were observed in two independent samples of patients affected by schizophrenia compared to healthy controls when global signal was not regressed out of the data (Yang et al. 2014, 2017). Most important, in the work from Yang and collegues, such differences correlated with symptoms and were not present in other psychiatric conditions, suggesting that they are a diagnostically specific disease-related neural phenotype.

In the present work we identified the three most recurrent patterns of activation during the resting state and investigated how such metastable substates differently alternate to recreate the healthy and pathological state, defined as the corresponding PMS. In line with results from Yang and collegues commented above, we found that the SCZ group showed a significantly increased probability of occurrence of a substate characterised by widespread synchronisation of all nodes. This means that the brain activity dynamics in the pathological condition are most frequently dominated by the global signal, defined as covariations averaged across all brain ROIs, as compared to the healthy population. This difference was specific for the SCZ condition and was not present in other psychiatric conditions, like bipolar disorder and ADHD.

A rise in the global signal indicates a tendency towards increased overall functional connectivity. It is interesting to notice that, in our data, the increased occurrence of the global state does not correspond to a significant increase in global static functional connectivity. This would indicate that the brain falls more often into a hyperconnected state rather than being generally hyperconnected at all times.

Increased connectivity might seem contradictory with the dysconnectivity hypothesis used to describe schizophrenia. Connectome investigations are in fact generally concordant in showing a widespread reduction of long-range structural connections in this disease and disappearance of network connector hubs (Cabral, Kringelbach, and Deco 2012; Van Den Heuvel et al. 2013; Mastrandrea et al. 2021; Rubinov and Bullmore 2013). However, the way functional alterations emerge from structural changes is complex (Cabral 2013) and not fully clear, and results regarding functional connectivity in schizophrenia are more contradictory, with some works indicating a tendency to reduced connectivity (Don 2018, Li 2019) and others showing evidence of increased connectivity (Driesen et al. 2013; Jafri et al. 2008; Yang et al. 2014). As discussed by Fornito and Bullmore such a dissociation could be explained as a compensatory process, as differentiation of neural activity characterized by a breakdown of normally segregated neural activity, or as mixture of both (Fornito and Bullmore 2015). Moreover, an increase in functional connectivity in schizophrenia is coherent with many lines of research suggesting that this disorder may involve extensive disturbances in the NMDA glutamate receptor, and a resulting altered balance of excitation and inhibition (Marín 2012). A generalised local disinhibition might in fact lead to generalised hyperconnectivity (Yang et al. 2016).

From the technical point of view, it is important to notice that GSR can alter the distribution of functional connectivity estimates in the data, leading to the emergence of spurious connectivity values and the distortion of group differences (Saad et al. 2012). GSR has also been shown to invert the directionality of specific functional alterations in schizophrenia (Yang et al. 2014). The authors proved in fact that, while data showed decreased functional connectivity before GSR it can show increased connectivity after correction (Fornito and Bullmore 2015). Also, when looking at functional connectivity of specific networks, the confusion can emerge from the fact that the activity of specific areas is differently influenced by global signal, and therefore by its removal (Yang et al. 2017).

The current work stands in favour of more conservative pre-processing procedure that does not get rid of global signal when analysing psychotic disorders, as it contains patho-physiologically relevant information. Moreover, LEiDA analysis allows one to separately investigate differences in global signal dynamics and alterations in specific connectivity patterns. This way it adds new evidence on the presence of a generalised increase in dynamical functional connectivity in SCZ patients when GS is not removed, while showing at the same time how frontal areas are less recruited in the pathology, as discussed in the following section.

### Schizophrenia is characterised by alteration in specific patterns of functional brain organisation

In our study, the second metastable substate was found to occur less frequently in the SCZ group as compared to the HC condition. This substate represents a pattern of activation where several areas from the medial prefrontal cortex synchronise between each other and with the parahippocampal cortex, hippocampus and nucleus accumbens, while desynchronising from the rest of the brain. When compared with the most known Yeo networks (Yeo et al. 2011), this connectivity pattern mostly correlates with the default mode and the limbic networks, both proven to be crucially involved in schizophrenia (Hu et al. 2017). As such, this substate is generally characterised by the synchronisation of high-order associative areas involved in memory, integration, and top-down regulation. All these areas have been proven to be altered in schizophrenia, in particular areas within the prefrontal cortex and the anterior-cingulate (Bersani et al. 2014; Rolls et al. 2008; Wang et al. 2015; Zhou et al. 2005). Moreover, differences in the frequency of recruitment of such network of areas is in line with the view of schizophrenia as a pathology determined by an impaired top-down regulation (Sun et al. 2021).

Notably, differences found here between the healthy and the pathological condition could not be explained by differences in static measures of functional connectivity or strength, proving the importance of integrating those measurements with dynamical analysis that do take into account the dimension of time.

### Towards the identification of vulnerable areas in the healthy brain model

We provided evidence for a critical role of the malfunction of single brain regions to promote the emergence of pathological-like dynamics. To do so we effectively reproduced the above described dynamical properties of the healthy condition data using an in-silico whole-brain model, by adjusting local and global parameters until optimal fitting of the empirical PMS space was reached. Then we systematically perturbed each pair of nodes by altering the local bifurcation parameter and we confronted the resulting dynamics with the empirical PMS of patients affected by schizophrenia. We found that it was possible to affect global dynamical patterns to induce a shift toward the pathological condition by shifting the local parameter *a* of the network toward positive values.

Notably, modifying the local properties in such way means tuning up the level of internal synchronisation and increasing the local intrinsic activity, potentially reflecting a E/I elevation that has been shown to result into increased functional connectivity (Yang et al. 2016). We can notice that, as a result of effective perturbation, we observe an increase of the probability of occurrence of the first metastable substate, representing dominance of the global signal over whole brain dynamics. In favour of this interpretation, we also showed that global FC increased as a result of the perturbation. This result is coherent with the increased level of global signal found at the empirical level in the pathological condition.

Specifically, we identified vulnerable areas in parieto-temporal regions involved in primary and higher-order sensory processing, such as the postcentral cortex and fusiform cortex, as well as associative areas involved in the processing and integration of external or internal incoming stimuli, such as the insula, inferior parietal, and middle temporal cortexes, and in memory, such as the hippocampus. This result would indicate that an increased engagement of those areas alters global dynamics, resulting in a pathological-like resting state, in line with other work highlighting dysfunctional alterations in sensory processing and integration in schizophrenia (Dong et al. 2019; Zhang, Guo, and Tian 2019). Increased intrinsic synchronisation of superior frontal cortex and anterior cingulate has been associated with delusions (Gao et al. 2015) and emergence of hallucinations has been related to increased intrinsic activity in sensory cortexes (visual-pericalcarine, somatosensory-postcentral), associative cortexes for integration and self-representation (insula) and for memory, (medial temporal lobe and hippocampus) and areas involved in language production (Broca)(Zmigrod et al. 2016). One recent cross-sectional study (Griffa et al. 2019) investigated the role of a set of critical areas in characterising the progressive emergence of the connectome disruption in psychosis. Note that many of the areas highlighted in that study, for example superior frontal and inferior parietal cortex, coincide with those emerged as vulnerable in our current work, evidencing their crucial role in the pathology.

### Towards the identification of areas sensitive to perturbation in the pathological brain model

We provided evidence that is also possible to induce a shift in the opposite direction to simulate healthy-like brain dynamics by perturbing individual pair of areas in the model of the pathological condition. In this way, we identified those areas that could be involved in potential compensation to restore a correct functioning in schizophrenia.

We found that such a shift toward the healthy condition can be induced by moving the local bifurcation parameter toward more negative values, and therefore reducing rhythmic oscillations (noisy regime). Note that, as shown by previous research (López-González et al. 2021), nodes that perform as functional hubs of connectivity in the healthy condition are associated with more negative values of *a* compared to the rest of the nodes, which preferentially set at the verge of bifurcation (Deco et al. 2019). Hubs are defined as nodes which, thanks to their strategic connections, are important in facilitating integration by spreading information between different brain regions.

In this work we showed that assigning negative values of *a* to specific regions significantly alters the global dynamics of the whole system, restoring healthy dynamics. Associative and cognitive regions were especially responsive to this kind of perturbation. In particular, we identified as critical specific prefrontal, temporal, and parietal associative and limbic areas, such as the orbitofrontal cortex, the cingulate, lingual cortex, IFG (pars orbitalis) and inferior temporal cortex, that are known to be relevant for schizophrenia and could represent impaired hubs. Note that disruption of associative and limbic areas, mainly in prefrontal but also in parietal and temporal regions, has been repeatedly found in many studies, as reviewed in Rubinov & Bullmore, 2013, and the consequent loss of hierarchical and networks properties has been proposed as one of the possible underlying mechanisms of schizophrenia (Bassett et al. 2008; Cabral et al. 2012; Van Den Heuvel et al. 2013). In the current work we have shown that the perturbation of one of those nodes alters not only the strength of connection of the perturbed area with the rest of the brain, but also affects the levels of global functional connectivity in the system. For example, a perturbation in the medial orbitofrontal cortex induces a more complex combination of reductions and increments of pairwise FC, indicating how local and global dynamics are interconnected. Accordingly, computational models have confirmed that alteration of hub nodes yields to a complex pattern of increases and decreases in functional connectivity (Alstott et al. 2009). Perturbation of a few areas - the superior temporal sulcus, the IFG (pars triangularis) and the pallidum-induces a shift in both directions, from the healthy to the pathological state and vice versa. These areas have been shown to be involved in psychiatric conditions, and in particular in psychotic disorders. For example, the IFG (pars triangularis) has been shown to be associated with the severity of symptoms typical of psychosis, such as thought disorder and delusion (Jung et al. 2019). Similarly, alterations in the superior temporal sulcus have been proved to correlate negatively with severity of hallucinations and thought disorder (Rajarethinam et al. 2000).

### LIMITATIONS AND FUTURE APPLICATIONS

To the best of our knowledge, this work is a first demonstration of the possibilities offered by the application of in silico perturbational approach to the study of schizophrenia. One limitation of this work is the use of the same standard structural connectivity template for both the control and the pathological conditions. Building both models using a common connectome allowed us to meaningfully compare global coupling and functional dynamics changes between conditions, which was the aim of the work. This way, though, it was not possible to evaluate the effect of local structural alterations. Future research should focus on integrating such features in the models. A second limitation is represented by the sample size. In fact, despite being in line with similar works in the field, the size of the dataset analysed in the current work is relatively low given the wide heterogeneity of the disease (Marek et al. 2022). For this reason, the current findings will need to be replicated in larger datasets, to verify their generalizability. In order to move to a more individualised approach and address heterogeneity limitations, it would be also extremely important to switch from group-level to patient-level analysis and simulations. Finally, future research should apply multimodal imaging such as EEG-fMRI or in OPM-MEG in order to fully solve the debate on the extent to which global signal reflects meaningful neuronal processes (Glasser 2019, Zhang and Northoff 2022) and whether a portion of it should be attributed to artefacts (Power 2019, Aquino 2022).

## CONCLUSIONS

This study uses dynamical analysis to separately investigate alterations in global signal from other relevant altered brain dynamics in schizophrenia. Results show a significantly increased probability of occurrence of global signal in schizophrenia that is not present in other pathologies, and a decreased recruitment of internally driven networks. Furthermore, in this study in silico perturbation is used to identify critical areas for the pathology, potentially opening the way to new clinical investigations.

## ACKNOWLEDGEMENTS

The project that gave rise to these results received the support of a fellowship from “la Caixa” foundation “(ID 100010434)”. The fellowship code is: “(LCF/BQ/DI19/11730048)”, and financed L.M work. In addition, G.D., M.V and L.M. were supported by the Human Brain Project Specific Grant Agreement 3 Grant agreement no. “(945539)” and by the Spanish Research Project AWAKENING: using whole-brain models perturbational approaches for predicting external stimulation to force transitions between different brain states, ref. “(PID2019-105772GB-I00/AEI/10.13039/501100011033)”, financed by the Spanish Ministry of Science, Innovation and Universities (MCIU), State Research Agency (AEI). G.D. and M.V. were also supported by the project “Clúster Emergent del Cervell Humà” “(CECH, ref. 001-P-001682)”, within the framework of the European Research Development Fund Operational Program of Catalonia 2014–2020. M.L.K. is supported by the Center for Music in the Brain, funded by the Danish National Research Foundation “(DNRF117)”, and Centre for Eudaimonia and Human Flourishing at Linacre College funded by the Pettit and Carlsberg Foundations.

## AUTHOR CONTRIBUTIONS

L.M, G.D., M.V.V. and M.L.K designed the research; K.A and A.F. pre-processed the data; L.M., M.V.V. and C.K. analysed the data; L.M. and M.V.V. wrote the manuscript; M.V.V and G.D. supervised the research. All authors contributed to the editing of the manuscript. Correspondent author: L.M.

