## Supplementary data for "Using in silico perturbational approach to identify critical areas in schizophrenia"

**SUPPLEMENTARY MATERIAL**

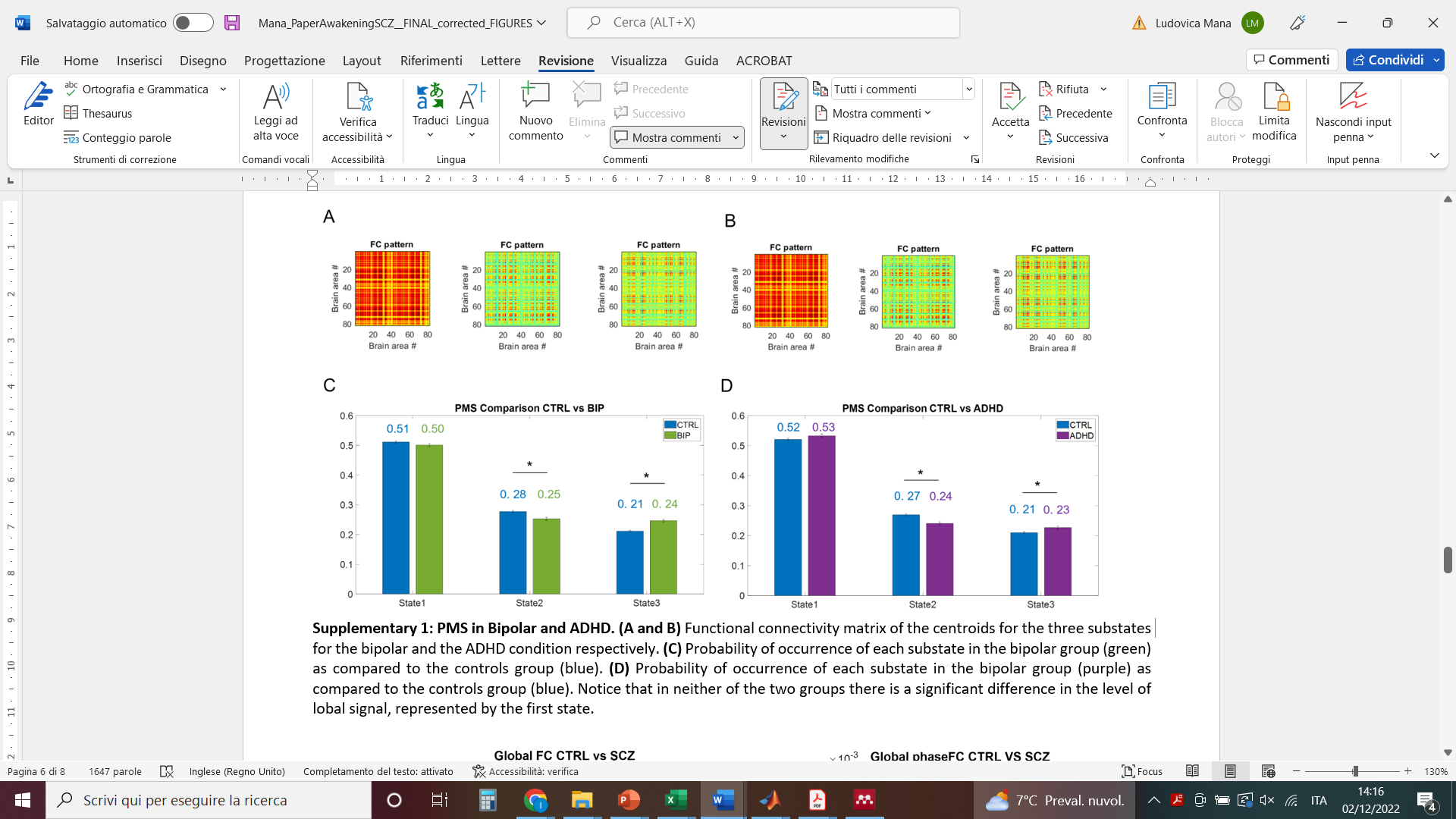

***Supplementary 1: PMS in Bipolar and ADHD. (A and B)*** *Functional connectivity matrix of the centroids for the three substates for the bipolar and the ADHD condition respectively.* ***(C)*** *Probability of occurrence of each substate in the bipolar group (green) as compared to the controls group (blue).* ***(D)*** *Probability of occurrence of each substate in the bipolar group (purple) as compared to the controls group (blue). Notice that in neither of the two groups there is a significant difference in the level of global signal, represented by the first state.*

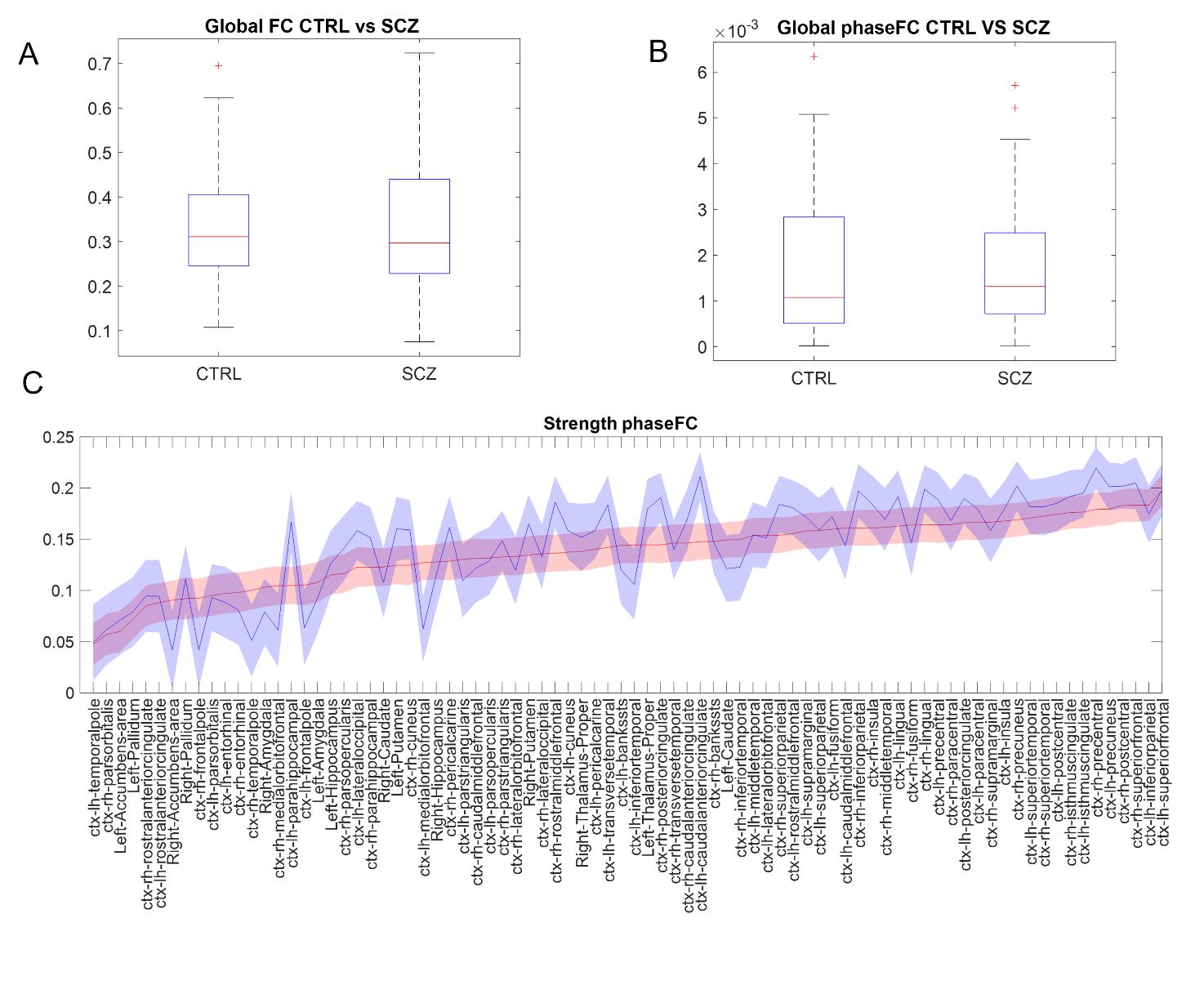

***Supplementary 2: Static analysis.*** *No significant differences were found between the levels of global static functional connectivity* ***(A)*** *and global phase coherence****(B)*** *of the schizophrenic group as compared to the healthy controls, nor for the levels of strength of any brain areas* ***(C)****.*

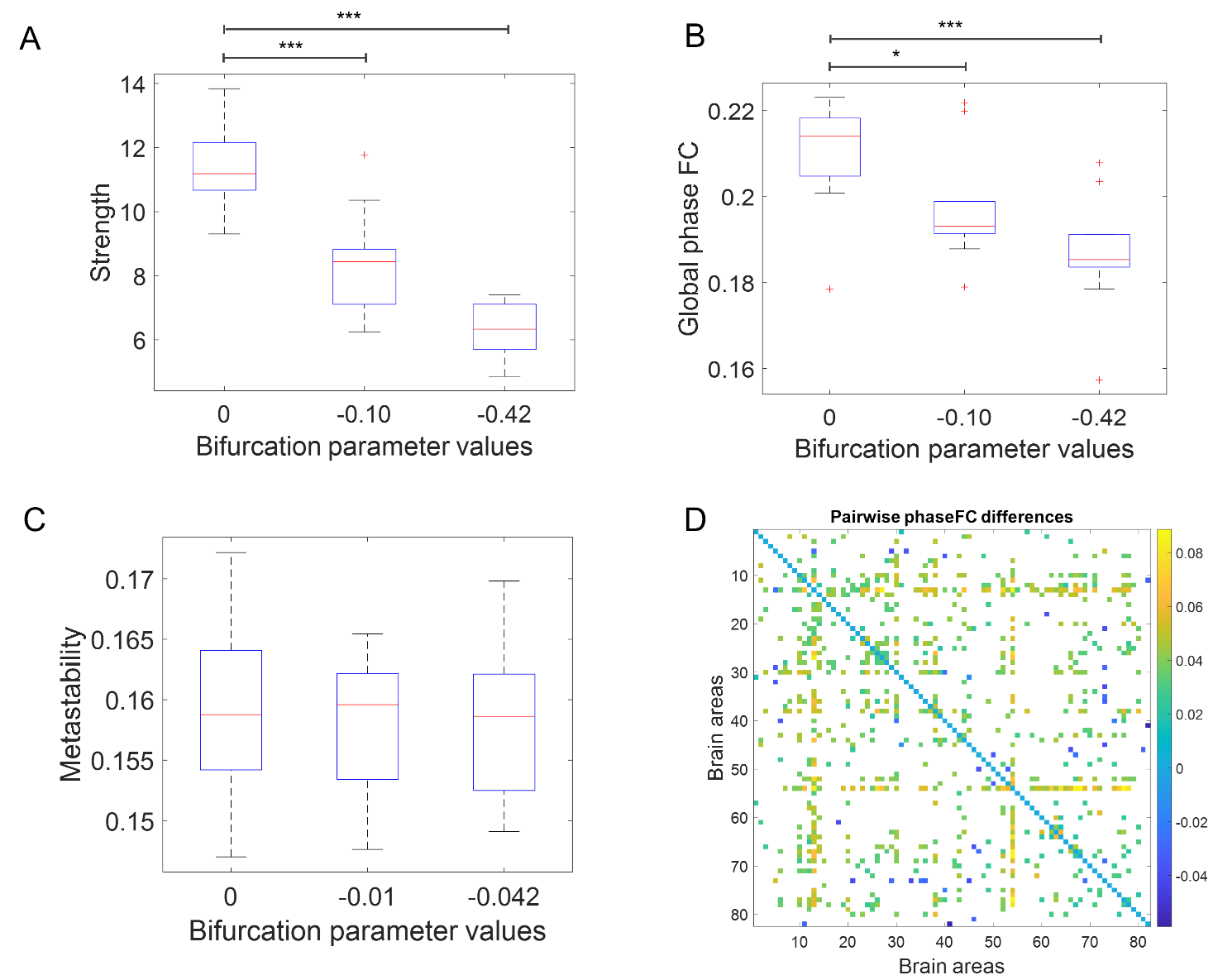

**Supplementary 3: Effect of perturbation on functional connectivity.** Analysis of effective perturbation on medial orbitofrontal cortex, inducing a shift from pathological to healthy state by moving the parameter toward negative values. **(A)** Significant progressive reduction of the strenght of the perturbed area at different values of perturbation. **(B)** Significant progressive reduction of the global phase coherence as a result of the perturbation of medial orbitofrontal cortex at different values of perturbation. **(C)** Metastability does not significantly change as an effect of perturbation. **(D)** Pairwise differences in phase coherence as an effect of perturbation (a=0 – a= -0.105). A mask is applied to only show values significantly affected (Wilcoxon ranksum test, p<0,05 prior to multiple comparison correction). We can notice a prevalent tendency toward decreased connectivity (positive difference between non perturbed and perturbed values), with some pairs of nodes showing increased connectivity (negative difference between non perturbed and perturbed values). No value survive correction for multiple comparisons.

***Table1: Vulnerable areas	Table2: sensitive areas***

Banks of superior temporal sulcus

Caudal anterior cingulate cortex

Inferior temporal cortex

Isthmus cingulate cortex

Lateral orbitofrontal cortex

Lingual cortex

Medial orbitofrontal cortex

Parahippocampus

IFG (Pars orbitalis)

IFG (Pars triangularis)

Posterior cingulate cortex

Precentral cortex

Caudate

Pallidum

Banks of superior temporal sulcus

Fusiform cortex

Inferior parietal cortex (IFG)

Middle temporal cortex

IFG (Pars triangularis)

IFG (Post central cortex)

Rostral anterior cingulate cortex

Superior Frontal cortex

Supra marginal cortex

Insula

Pallidum

Hippocampus
